# Quantifying Unbiased Conformational Ensembles from Biased Simulations Using ShapeGMM

**DOI:** 10.1101/2024.01.22.576692

**Authors:** Subarna Sasmal, Triasha Pal, Glen M. Hocky, Martin McCullagh

## Abstract

Quantifying the conformational ensembles of biomolecules is fundamental to describing mechanisms of processes such as ligand binding and allosteric regulation. Accurate quantification of these ensembles remains a challenge for all but the simplest molecules. One such challenge is insufficient sampling which enhanced sampling approaches, such as metadynamics, were designed to overcome; however, the non-uniform frame weights that result from many of these approaches present an additional challenge to ensemble quantification techniques such as Markov State Modeling or structural clustering. Here, we present rigorous inclusion of non-uniform frame weights into a structural clustering method entitled shapeGMM. The shapeGMM method fits a Gaussian mixture model to particle positions, and here we advance that approach by incorporating nonuniform frame weights in the estimates of all parameters of the model. The resulting models are high dimensional probability densities for the unbiased systems from which we can compute important thermodynamic properties such as relative free energies and configurational entropy. The accuracy of this approach is demonstrated by the quantitative agreement between GMMs computed by Hamiltonian reweighting and direct simulation of a coarse-grained helix model system. Furthermore, the relative free energy computed from a high dimensional probability density of alanine dipeptide reweighted from a metadynamics simulation quantitatively reproduces the metadynamics free energy in the basins. Finally, the method identifies hidden structures along the actin globular to filamentous-like structural transition from a metadynamics simulation on a linear discriminant analysis coordinate trained on GMM states, demonstrating the broad applicability of combining our prior and new methods, and illustrating how structural clustering of biased data can lead to biophysical insight. Combined, these results demonstrate that frame-weighted shapeGMM is a powerful approach to quantify biomolecular ensembles from biased simulations.

---

Conformational ensembles of molecules dictate their thermodynamic properties. Conventional molecular dynamics (MD) simulations allow us to sample these ensembles but suffer from the so-called rare event problem. A variety of enhanced sampling techniques, such as Metadynamics (MetaD), ^1,2^ Adaptive Biasing Force, ^3^ Gaussian accelerated MD, ^4^ and Temperature Accelerated MD/Driven Adiabatic Free Energy Dynamics, ^5,6^ have been developed to promote faster sampling by effectively heating some degrees of freedom. Unfortunately, due to the biased sampling of many of these approaches, it is not obvious how to use the biased configurations in methods such as Markov State Models (MSMs) ^7,8^ and/or structural clustering approaches that quantify the conformational ensemble. Here, we adapt shapeGMM, ^9^ a probabilistic structural clustering method, to rigorously quantify the unbiased conformational ensembles generated from biased simulations. The result is a high dimensional Gaussian mixture model (GMM) characterizing the unbiased landscape that can be used to extract important thermodynamic quantities and to give additional insight beyond the low dimensional projections often used to represent free energy landscapes.

Meaningful quantification of conformational ensembles from large molecular simulations requires the grouping of similar frames using a clustering algorithm. Clustering algorithms for molecular simulation can be grouped into two categories: temporal and structural. Temporal clustering, such as spectral clustering of the transition matrix, ^10,11^ has been successfully applied to MD trajectories to achieve kinetically stable clusters for use in objects like MSMs. ^12–14^ Enhanced sampling techniques, however, can distort the underlying kinetics of the system making temporal clustering difficult to apply properly in these circumstances.

While there have been efforts to build MSMs from enhanced sampling data ^15,16^ it still remains a challenge. ^17^ Additionally, building MSMs relies on an initial structural clustering, making it critical to perform this step accurately even in the context of enhanced sampling. Structural clustering involves partitioning either frames or feature space into a finite number of elements. This can be achieved from enhanced sampling data but care must be taken to properly account for the non-uniform weights of the frames.

Previous efforts to use structural clustering algorithms on enhanced sampling simulations have focused on deterministic, as opposed to probabilistic, algorithms. The distinction between these two approaches lies in data partitioning. Deterministic techniques use a hard assignment while probabilistic ones allows for a soft assignment. In a deterministic approach, all that matters is the cluster populations and, once the data has been clustered, these populations can be reweighted based on enhanced sampling frame weights. ^15,18^ Probabilistic clustering is an appealing approach for MD data because the resulting probabilities have strong theoretical ties to thermodynamic quantities. Reweighting the cluster populations is not satisfactory for probabilistic methods such as GMMs, as the frame weights will affect the means and covariances of each cluster. It is possible to use multiple copies of frames to approximately account for the frame weights but this can yield intractably large trajectories and inaccuracies due to rounding.

In this work, we present an adaptation to shapeGMM, ^9^ a probabilistic structural clustering method on particle positions, to directly account for non-uniform frame weights. As opposed to introducing copies of input data and maintaining uniform weights, the current method directly accounts for non-uniform frame weights and is thus more efficient and scalable than the alternative. In the next section we briefly introduce the shapeGMM method and the adaptations necessary to account for non-uniform frame weights. This is followed by a demonstration of the method on three examples of increasing difficulty, specifically demonstrating that our intuitive choices of frame weights from Metadynamics simulations result in a reliable clustering procedure. We show in benchmark cases how this method can yield thermodynamic quantities directly, and use the complex case of actin flattening to show how a weighted shapeGMM can give physical insight into the conformations sampled, in a case where unbiased simulation would not be a practical option. In addition, frame-weighted shapeGMM is implemented in an easy-to-use python package (pip install shapeGMMTorch).

## Theory and Methods

### Overview of shapeGMM

In shapeGMM, a particular configuration of a macromolecule is represented by a particle position matrix, ***x***_*i*_, of order *N* × 3, where *N* is the number of particles being considered for clustering. To account for translational and rotational invariance, the proper feature for clustering purposes is an equivalence class,

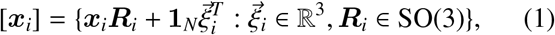

Where 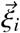 is a translation in ℝ^3^, ***R***_*i*_ is a rotation ℝ^3^ ℝ^3^, and **1**_*N*_ is the *N*× 1 vector of ones. [***x***_*i*_] is thus the set of all rigid body transformations, or orbit, of ***x***_*i*_.

The shapeGMM probability density is a Gaussian mixture given by

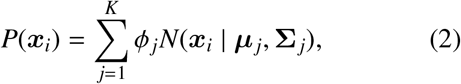

where the sum is over the *K* Gaussian mixture components, *ϕ* _*j*_ is the weight of component *j*, and *N*(***x***_*i*_ | ***µ*** _*j*_,Σ _*j*_) is a normalized multivariate Gaussian given by

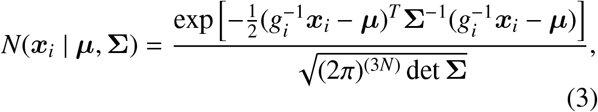

where ***µ*** is the mean or minimum energy structure,Σ is the covariance, and 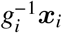 is the element of the equivalence class, [***x***_*i*_], that minimizes the squared Mahalan-bonis distance in the argument of the exponent. Determining the proper transformation, *g*_*i*_, is achieved by translating all frames to the origin and then determining an optimal rotation matrix. Cartesian and quaternion-based algorithms for determining optimal rotation matrices are known for two forms of the covariance were considered Σ ∝ ***I***_3*N*_ ^19,20^ or Σ = Σ_*N*_⊗ ***I***_3_, ^21,22^ where Σ_*N*_ is the *N*×*N* covariance matrix and ⊗ denotes a Kronecker product. In this manuscript, we employ only the more general Kronecker product covariance.

### Incorporating Non-uniform Frame Weights in shapeGMM

Previously, each frame in ShapeGMM was considered to be equally weighted. Approximate weighting of frames could be taken into account by including frames multiple times in the training data to give them more importance, however this introduces the imprecision of rounding to the nearest integer and can be extremely computationally expensive due to the large increase in amount of training data. Here, we take non-uniform frame weights into account by performing weighted averages in the Expectation Maximization estimate of model parameters 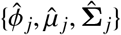, consistent with other fixed-weight GMM procedures. ^23^ Considering a normalized set of frame weights, {*w*_*i*_} where 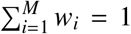 for *M* frames, their contribution to the probability can be accounted for by weighting the estimate of the posterior distribution of latent variables:

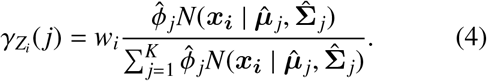

The frame weight will propagate to the estimate of component weights, means, and covariances in the Maximization step through 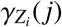:

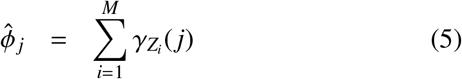

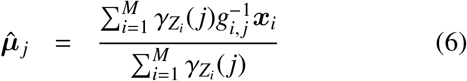

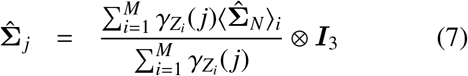

Additionally, the log likelihood per frame is computed as a weighted average

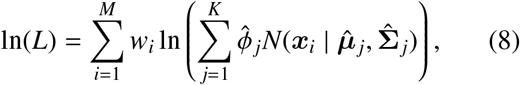

### Choosing Number of Clusters

Performing shapeGMM requires the user to choose a number of clusters, *K*. The “optimal” choice will be system and problem specific and potentially has no wrong answer. The choice is no different if you consider uniformly or non-uniformly weighted frames. We use a cluster scan with a combination of the elbow method and cross validation to assess if our choice of *K* is reasonable. A good choice of clusters based on this approach is to find the number of clusters where the increase in log-likelihood with *K* is decreasing fastest, which we can evaluate by choosing the minimum of the second derivative of ln(*L*) with respect to number of clusters. In practice, this works well for simple systems, but it may be hard to pick a “best” choice for more complex systems, so we may seek a choice that is physically interpretable.

### Implementation

We have completely rewritten shapeGMM in pyTorch for computational efficiency and ability to use GPUs. The current implementation takes an array of frame weights as an optional argument to both the fit and predict functions (the code defaults to uniform weights). The pyTorch implementation is significantly faster than the original version and is available both on github (https://github.com/mccullaghlab/shapeGMMTorch) and PyPI (pip install shapeGMMTorch). Examples are also provided on that github, and all examples from this paper are provided in a second github page discussed below.

### Choosing Training Sets

For non-uniformly weighted frames the choice of training set may be important. We have attempted a variety of training set sampling schemes and have found that, at least for the frame weight distributions that we have encountered, uniformly sampling the training data is at least as good as any importance sampling scheme. We discuss this further and show results for three different training set selection schemes for the beaded helix system in Sec. S1.

### Biasing and weighting frames

If configuration *x* is generated from an MD simulation at constant *T* and *V* then *P*(*x*) ∝ exp( −*H*(*x*)/*k*_*B*_*T*) where *H* is the system’s Hamiltonian. ^24^ If *x* is generated from an MD simulation at a different state point (e.g. different *T*) or with a different Hamiltonian, it is sampled from a different distribution *Q*(*x*). Samples from *Q* can be reweighted to *P* with weights ^24^

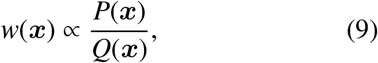

from which averages over *P* can be estimated. This approach is only effective if *Q* and *P* are finite over the same domain. Nonetheless, Eq. 9 underlies many enhanced sampling approaches, for example, it is the basis of the original formulation of umbrella sampling. ^25^ By including weights in shapeGMM, we can predict the importance of clusters at nearby state-points or for similar systems.

### Thermodynamic Quantities from ShapeGMM

Many Thermodynamic quantities can be computed from fit shapeGMM probability densities. One such quantity is the configurational entropy,

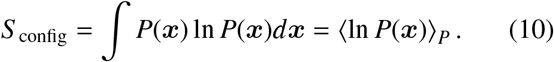

The configurational entropy has an analytic solution for a single multivariate Gaussian but for the general mixture of multivariate Gaussians we use sampling and Monte Carlo integration to approximate the integral. To do so accurately requires that we generate points from the shapeGMM objects and not just use the trajectory on which the object was fit. We have introduced a generate function as an attribute to a fit shapeGMM object that produces configurations sampled from the underlying trained distribution.

The second Thermodynamic quantity we consider is the free energy cost to move from one distribution to another. This is also known as the relative entropy or Kullback-Leibler divergence and the cost to go from distribution *Q* to distribution *P* is given by

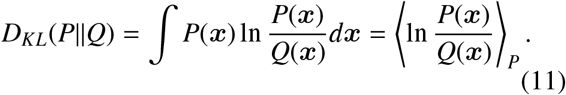

Here, again, we generate points from distribution *P* and average the difference in log likelihoods of these points in *P* and *Q* to assess this value. It should be noted that this is a non-equilibrium free energy and is thus not necessarily symmetric. ^26,27^ The quantity can prove useful in applications, for example measuring the free energy cost to shift a distribution from an apo to a ligand-bound state, for example. ^28,29^

A symmetric metric is useful when comparing the similarity of two distributions. Here we opt for the Jensen-Shannon divergence (*JS D*) ^30^ given by

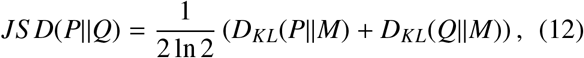

where 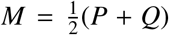 is the midpoint distribution between *P* and *Q. JS D* is restricted to be between 0 and 1.

All three of these measures have been implemented in the similarities library of the shapeGMM code. They use point generation and Monte Carlo sampling to assess the integrals and thus return both the mean value and the standard error.

## Results and Discussion

### Proof of Concept: Reweighting the Beaded Helix

To demonstrate the accuracy of the frame-weighted shapeGMM process we perform Hamiltonian reweighting of a non-harmonic beaded helix previously studied in Refs. 9,31. The system is composed of 12 beads connected in sequential fashion by stiff harmonic bonds. Every fifth pairwise interaction is given by an attractive Lennard-Jones potential with well depth *ϵ*. The value of *ϵ* relative to *kT* dictates the stability of an alpha-helix-like structure as compared to a completely disordered state. Additionally, because of the symmetry of the model both the left- and right-handed helices have equal probability no matter the value of *ϵ*. A value of *ϵ* = 6 in reduced units forms stable helices while allowing transitions between the two folded states; here we performed a long unbiased trajectory to sample both left and right states, as well as possibly intermediates (see Sec. A1 for details).

Fig. 1A shows the unbiased free energy for this system using *ϵ* = 6 computed as *F*(*s*) = −ln *P*(*s*) for a linear discriminant (LD) reaction coordinate, ^32^ shown as a black dashed line. By performing a scan over number of clusters on 100k frames from an unbiased trajectory, we identify three clusters as the optimal number by observing a definite kink in the curves in Fig. 1B and the presence of a minimum in the second derivative in Fig. 1C. These clusters correspond to the left- and right-helical states as well as a partially unfolded intermediate cluster, examples shown in Fig. 1A.

**Figure 1:**
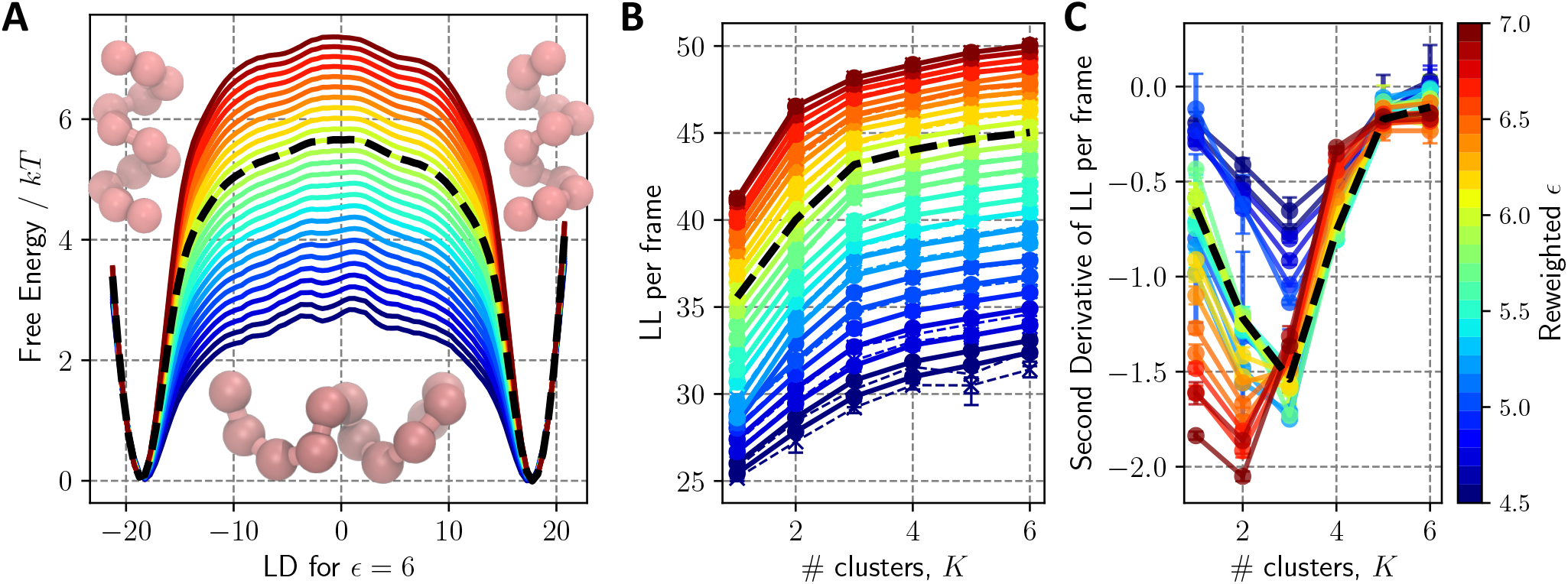
Beaded helix *ϵ* reweighting. Trajectory data for a 12 bead polymer having *i, i* + 4 interactions with strength *ϵ* = 6 was reweighted to predict the ensemble for *ϵ* values ranging from 4.5 to 7 in increments of 0.1. (A) The corresponding free energies as a function of the linear discriminant (LD) between the two helices are plotted with *ϵ* values denoted by the color bar on the right-hand. The weights per frame were fed in to shapeGMM to perform a cluster scan. (B) The resulting log likelihood per frame as a function of number of clusters from the cluster scan. (C) Second derivative of the curves from B. Error bars in (B) and (C) are estimated as the standard deviation from three different training sets. The true curve for *ϵ* = 6 is given in black dashed lines in all three panels.

Reweighted clustering of the beaded helix system predicts that the prevalence of the partially unfolded intermediate will disappear by *ϵ* = 6.5. To demonstrate this, we performed *frame-weighted* shapeGMM cluster scans of our trajectory at *ϵ* = 6 with weights corresponding to *ϵ* values ranging from 4.5 to 7.0 in increments of 0.1. Given that the samples come from a Boltzmann distribution, the weights for each frame given by Eq. 9 are 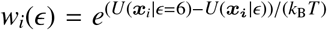. The log likelihood of the shapeGMM fits as a function of number of clusters are shown in Fig. 1B,C with *ϵ* values indicated in the color bar on the right. We see that as *ϵ* increases from 6, the minimum in the second derivative moves from 3 clusters to 2 cluster. The transition happens between *ϵ* = 6.4 and *ϵ* = 6.5. This suggests that a simulation run at *ϵ* values of greater than 6.4 (in reduced units) will not exhibit the partially unfolded third cluster. These results are consistent with the increasing free energy barrier height as a function of *ϵ* depicted in Fig. 1A.

The reweighting of *ϵ* for the beaded helix example also predicts that only one cluster will be present for small *ϵ*. In Fig. 1B, the elbow at 3 clusters is evident for *ϵ* values as low as *ϵ* = 5 and becomes less pronounced below this threshold. While a minimum at 3 clusters is still observed in the second derivative plot for *ϵ* = 4.5, the trend is clear that as *ϵ* becomes small the choice of anything other than 1 cluster is less well supported by the elbow heuristic. This is an expected result, and consistent with the reduced free-energy barriers observed for small *ϵ* in Fig. 1A, as *ϵ* approach thermal energy the prevalence of anything other than an unfolded state is entropically unfavorable.

ShapeGMM reweighted clustering also produces *quantitatively accurate* probability densities for the the beaded helix. To demonstrate this, we compute a reweighted shapeGMM object (ϵ = 6 → 8) to a shapeGMM object trained on an unbiased trajectory at *ϵ* = 8, which we refer to as ground truth (GT). Because, as predicted, transitions at *ϵ* = 8 are very unlikely, this object is trained on simulations, each with 100k frames, initiated from left and right helices and concatenated. Two controls are included that are fit to the *ϵ* = 6 trajectory without reweighting: the predicted 3 cluster object and a 2 cluster object in which we remove the partially unfolded cluster. To quantitatively compare between two probability densities we use two similarity metrics, both described above in more detail and introduced as Eqs. 10,11: Jensen-Shannon divergence (JSD) and change in configurational entropy *S* _config_. These similarity metrics between the GT and the three different shapeGMM objects are tabulated in Tbl. 1. JSD is a symmetric metric bounded between 0 and 1 where 0 indicates no divergence and 1 indicates complete divergence between the two probabilities. The reweighted shapeGMM object demonstrates a very small JSD (0.0071 ± 0.0003) to the GT as compared to either of the *ϵ* = 6 objects (0.358 ± 0.002 and ± 0.401 0.002). This trend holds true when comparing relative *S*_config_’s with the difference in *S* _config_ between the reweighted and GT *ϵ* = 8 shapeGMM probabilities being within error of 0. These results indicate that the *ϵ* = 8 reweighted shapeGMM probability density is nearly identical to the GT.

**Table 1:** Similarity measures between three beaded helix probability densities fit from a simulation with *ϵ* = 6, *Q*, and the “ground-truth” (GT) probability density fit to a simulation at *ϵ* = 8. The reweigted probability densities are denoted by the number of clusters, *K*, and the value of *ϵ* used in reweighting, *ϵ*_*R*_. The three *Q*s are: *K* = 3 clusters and weighted to *ϵ*_*R*_ = 6.0, *K* = 2 clusters extracted from the three cluster *ϵ*_*R*_ = 6.0 model, an *K* = 2 clusters reweighted to *ϵ*_*R*_ = 8. The similarity measures are the Jensen-Shannon divergence (*JS D*) and the difference in configurational entropy 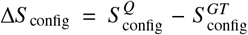. Error in the last digit is included in parentheses and are estimated as Monte Carlo sampling errors in estimating the integrals.

| $Q$ | | $JSD(GT\vert \vert Q)$ | $\Delta S_{\text{config}}/R$ |
| --- | --- | --- | --- |
| $K$ | $\epsilon_R$ | | |
| 3 | 6.0 | 0.401(2) | -7.22(3) |
| 2 | 6.0<sup>†</sup> | 0.358(2) | -4.02(2) |
| 2 | 8.0 | 0.0071(3) | 0.00(2) |

<sup>†</sup> The frames corresponding to the partially unfolded cluster were removed.

### Conformational States of Alanine Dipeptide from Metadynamics Simulations

Alanine Dipeptide (ADP) in vacuum is a common benchmark system for methods designed to sample and quantify conformational ensembles. In this work, we demonstrate that ADP MetaD simulations can be used directly to achieve equilibrium clustering using various estimates of the frame weights. In Well-tempered MetaD (WT-MetaD), a history dependent bias is generated by adding Gaussian hills to a grid at the current position in collective variable (CV) space ^2,33^ such that the bias at time *t* for CV value position ***s***_*i*_ is given by

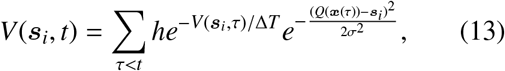

where *h* is Gaussian height, andΣ is the width, and *T* + Δ*T* is an effective sampling temperature for the CVs. Rather than setting Δ*T*, one typically chooses the bias factor γ = (*T* + Δ*T*)/*T*, which sets the smoothness of the sampled distribution. ^2,33^ Asymptotically, a free energy surface (FES) can be estimated from the applied bias by 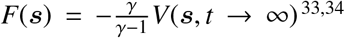 or using a reweighting scheme. ^33,35^ In MetaD, frames are generated from a time dependent Hamiltonian so the choice of frame weights for clustering is not obvious. Reweighting of MetaD trajectories to compute free energy surfaces has been accomplished through several different schemes.

For a static bias *V* added to the initial Hamiltonian, the weight of a frame given by Eq. 9 would be 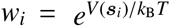. Our first choice of frame weights (termed ‘bias’) corresponds to using this formula even though the bias is time-dependent. A second choice that removes some of the time-dependence is to use 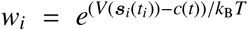, where 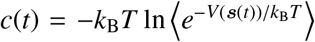 is the bias averaged over the CV grid at a fixed time. The quantity *V*(***s***_*i*_(*t*_*i*_)) − *c*(*t*) is called the “reweighting bias” and can be computed automatically in PLUMED, ^36^ hence we term clustering using this scheme ‘rbias.’ Finally, we evaluate another commonly used approach to compute Boltzmann weights of each frame post-facto, ^37^ which in the case of WT-MetaD would correspond to 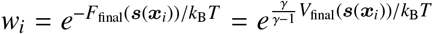 we label these weights ‘fbias’. Other more sophisticated reweighting schemes have also been proposed, e.g. in Refs. 37,38, but we did not test these here because, as will be seen, the bias, rbias, and fbias approaches all worked well for our test system. However, shapeGMM as implemented is capable of using any choice of frame weights.

For assessing the best choice of weights, we performed a 100 ns WT-MetaD simulation on ADP biasing backbone dihedral angles *ϕ* and *Ψ* using bias factor 10, saving every 1 ps to generate 100k frames (see Sec. A1 for full details). The five atoms involved in the *ϕ* and *Ψ* dihedral angles were chosen for shapeGMM clustering. The coordinates of these atoms and the frame weights from the four different schemes were fed into shapeGMM. The log likelihood per frame of the resulting fits as a function of number of clusters is shown in Fig. 2A. In general, the three non-uniformly weighted clustering objects result in significantly higher log likelihoods for equivalent numbers of clusters *K* > 2, indicating a much better fit to the underlying data. The significant kink in the cluster scans for the non-uniformly weighted objects at 2 clusters indicate that at least 2 clusters are necessary for a good fit to the data; there is still substantial increase going from 2 to 3 clusters, however, indicating that there may be additional insight gained at *K* = 3 and above, as we shall see.

**Figure 2:**
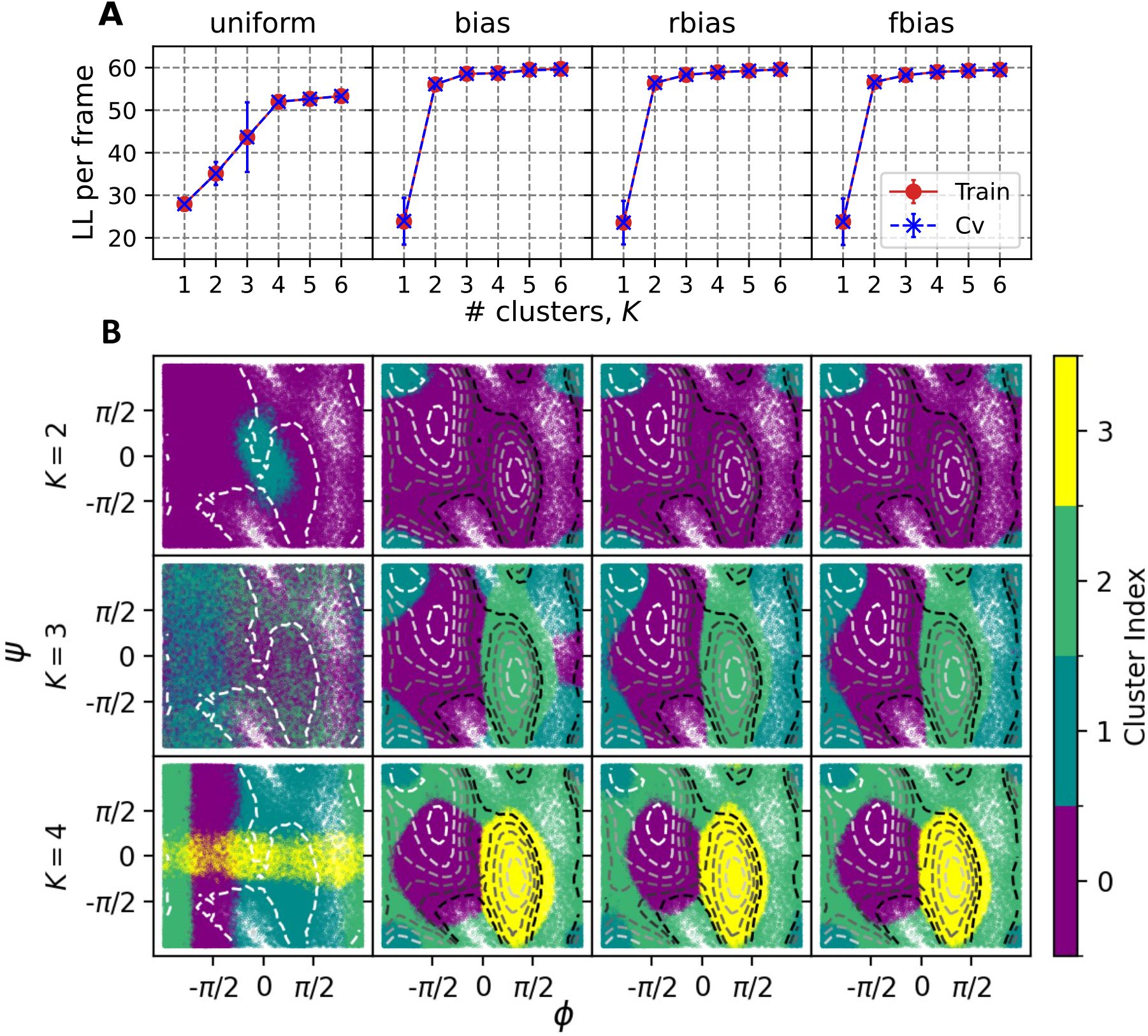
WT-MetaD simulation for ADP with BF 10. Each column represents a particular choice of weights been used in *frame-weighted* SGMM. (A) Cluster scans for each choice of frame weights using using 50k frames, 4 training sets and 10 attempts for each case. (B) Clusterings performed for *K* = 2, 3, 4 are shown by coloring each of 100K sampled points by their cluster assignment. Contour lines indicate the underlying free energy surface as computed from the WT-MetaD simulation via reweighting with the different choice of weights. Contours indicate free energy levels above the minimum from 1 to 11 kcal/mol with a spacing of 2 kcal/mol.

Non-uniform *frame-weighted* shapeGMM produces physically relevant clusterings. Fig. 2B indicates how sampled points are assigned to two, three, or four clusters when using each of the choices of frame weights, with the underlying free energy landscape computed from a weighted histogram using the same choice of weights as used for the clustering indicated by contour lines. Clustering with uniform weights has little correlation with the underlying free energy landscape, whereas performance is much better when using any of the non-uniform weighting schemes. Weighted clustering with *K* = 2 tends to split the landscape into one cluster covering the most extended upper-left “C5” basin near (−2,2), while using a second cluster to cover the rest of the landscape (see Ref. 39 for a naming convention). However, higher number of clusters allows for separating the upper left basin into its two constituent states, C5 and “C7eq” at (−2,1), while also revealing the presence of the minor “C7ax” state at (1,- 1). Slight differences in contour FES correspond with slight differences in the weighted cluster assignments; for example, in the *K* = 3 case the upper left and bottom left parts of the axial basin are disconnected at *Ψ* = 0 for bias weights but connected for rbias and fbias weights.

Non-uniform *frame-weighted* shapeGMM also works for standard (untempered) MetaD ^1,33^ with Δ*T* → ∞ For untempered MetaD, we favor rbias weights, because the final bias is not static and the instantaneous bias diverges, meaning that initial frames receive no weight. In Fig. S2, we show that shapeGMM clustering with rbias weights performs much better than equally weighted frames, and results are comparable to our study with WT-MetaD, indicating that *frame-weighted* shapeGMM can be extended to this method as well.

Non-uniform *frame-weighted* shapeGMM probability densities quantitatively capture the correct free energy basins. Because we know that the free energy in dihedral space is a good proxy for the configuration space of ADP, we here quantify the accuracy of our GMM fits (which are 15-dimensional objects) by predicting this FE landscape directly from the GMMs. To do so, we generate 1M samples in Cartesian space from each GMM object and compute the FES from an unweighted histogram of the backbone dihedral angles. Fig. 3 shows a comparison of these predicted FES with the reference FES computed directly from the WT-MetaD bias as described above. Here we see that uniform weights produces FES that span all of di-hedral space but whose minima are not centered on the true minima.

**Figure 3:**
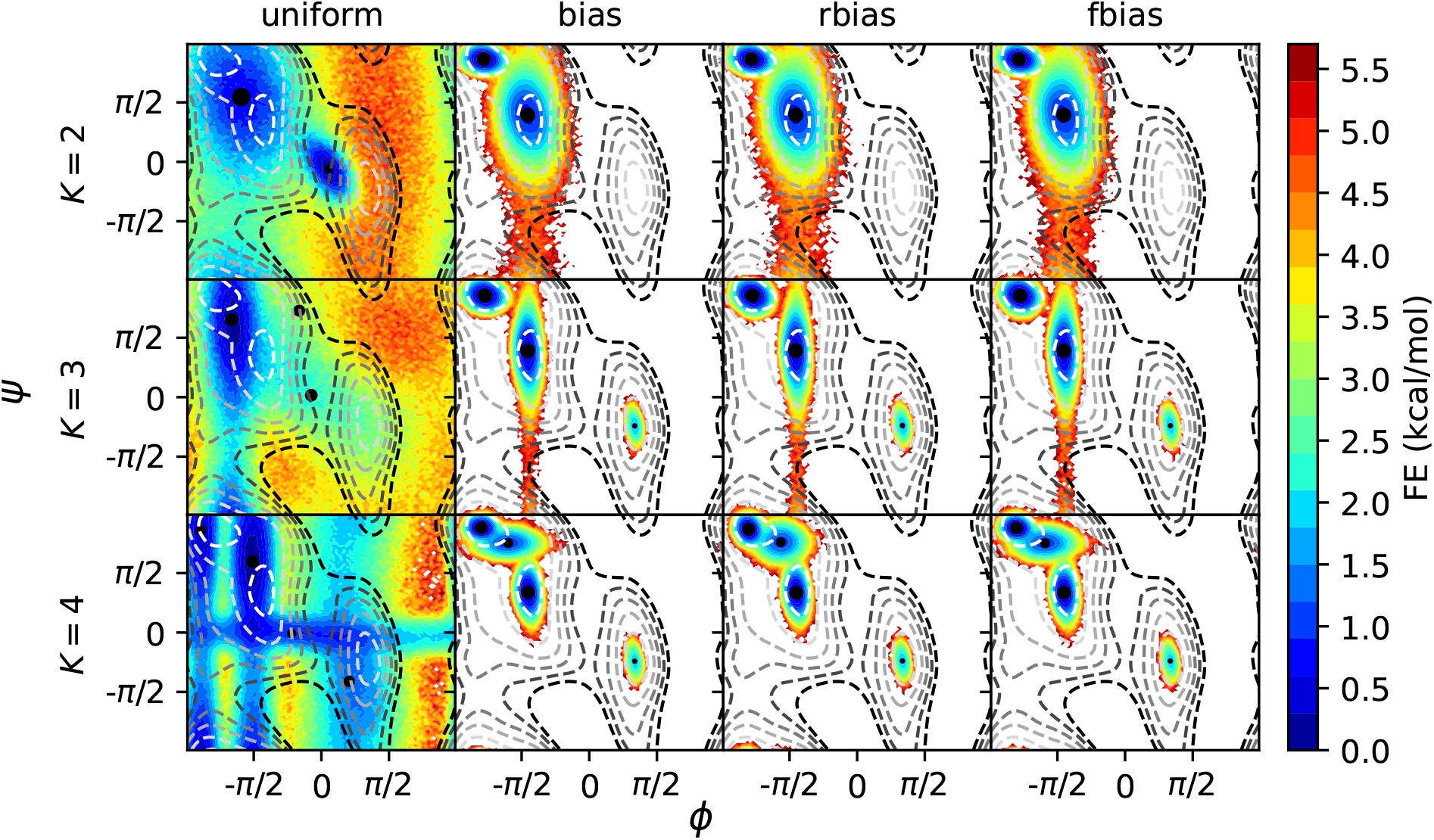
FE profiles obtained from GMM objects trained on BF=10 Metadynamics data. Each column corresponds to a different choice of bias and each row corresponds to a different number of clusters used. These are computed as unweighted histograms from 1M samples obtained from each GMM object. Black circles placed on the FEs are the centers calculated from the reference structures corresponding to different clusters, with the size indicating their relative population. Contour lines indicate the underlying free energy surface as computed from the WT-MetaD simulation, positioned at 1.0 to 11.0 kcal/mol with a spacing of 2 kcal/mol above the global minimum.

In contrast, the FES generated from the non-uniform weighting schemes demonstrates that the clustering above captures the nature of the underlying FES as well as could be expected given a limited number of clusters. FES for *K* = 2 capture the primary C7 equatorial global minimum and C5 metastable state, while going to three or more clusters also allows resolution of the minor C7 axial basin. As should be expected, the GMM objects only resolve the configurational landscape of our system around the minima, and cannot resolve (non-convex) high free energy regions. Importantly, we note that the results reflect an intrinsic error due to the fact that we are fitting an anharmonic landscape to a locally harmonic model, resulting in an over-estimate of the FES away from the minima. We can also compute a FES that covers the entire energy landscape using a Monte Carlo procedure described in Sec. S3, resulting in FES shown in Fig. S3 that are qualitatively correct but which also reflect the inherent overestimation of the Gaussian model.

The comparison of FESs can be further quantified by difference metrics which also provide an alternative metric to choose the best method or best number of clusters. In Fig. S4 we show both the root-mean-squared error (RMSE) for the sampled region and the JSD as compared to the reference FES. While the uniform weights perform poorly, we see that all other weights do comparably well for 3 or more clusters. Using RMSE as a metric, rbias weights are the most accurate by a small margin, and a five state clustering is the best within the range *K* = 2 to *K* = 6.

### Elucidating conformational states of the actin monomer

Up to this point, we have established that we can accurately train a GMM with data weighted from MetaD or Hamiltonian reweighting for small systems. In this section, we demonstrate that this approach can provide insight into data for a complex biochemical problem. The actin cytoskeleton, composed of filaments of actin, plays major roles in a wide range of active biological processes, including cell motility and division. ^41–43^ Actin filaments are non-covalent polymers that form from head-to-tail assembly of globular actin (G-actin), which is a 375-amino acid protein consisting of four primary subdomains (Fig. 4A). Each actin monomer contains a bound nucleotide that is in the form of ATP in G-actin and is eventually hydrolyzed to ADP as filaments “age”. ^42,44^ The polymerization from G-actin to filamentous actin (F-actin) results in a flattening of the protein which is characterized by a reduction of the *ϕ* dihedral angle shown in Fig. 4A. ^42^ An open question in the field is whether the flat state is metastable in solution, or whether it is only stabilized when contacting the end of a filament. ^45^ Additionally, the structural intermediates along the flattening pathway remain elusive.

**Figure 4:**
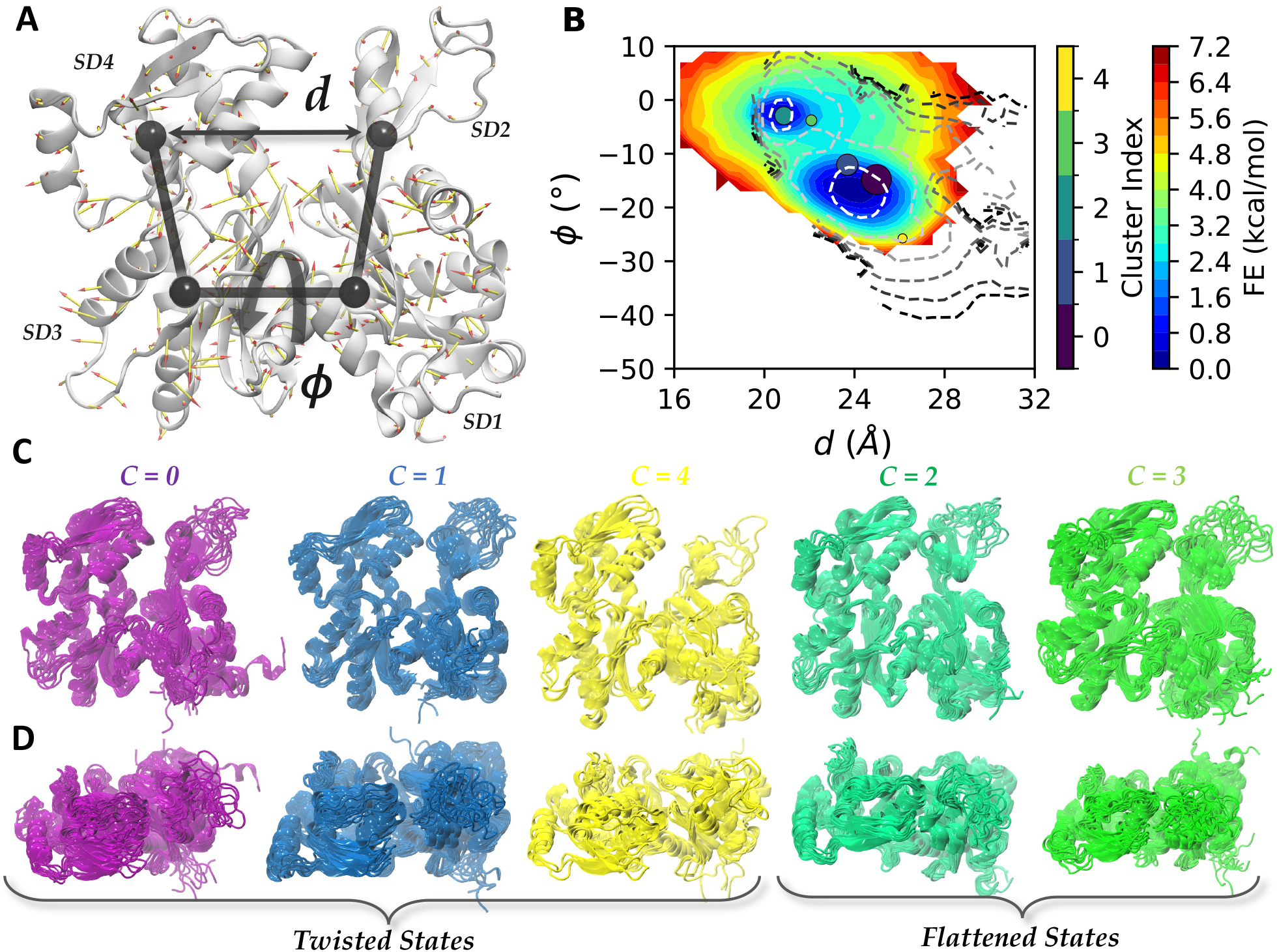
A. Cartoon representation of Actin monomer. The arrows representing the magnitude and directions of the LD vector acting on 375 *C*_α_ atoms. SD1 to SD4 are four subdomains defined for the monomer. ^40^ *d* is the distance between center of masses (COMs) of subdomains SD2 and SD4. *ϕ* is the dihedral angle defined using COMs of SD2-SD1-SD3-SD4 respectively B. FES calculated by performing an unweighted histogram of ∼ 1M samples generated from GMM. Contour lines represent the reweighted FE obtained from restarted OPES-MetaD trajectory using fbias frame weights. Contours are positioned at 1 to 11 kcal/mol with a spacing of 2 kcal/mol above the global minimum. Colored circles are the locations for different cluster centers weighted by relative population. C. Snapshots of frames belonging to different clusters (front view) D. Top view for the same.

Previous efforts to directly sample the flattening of G-actin have proven difficult. These efforts employed umbrella sampling or MetaD on two experimentally defined coordinates *ϕ* and *d* and demonstrate the difficulty in sampling the conformational landscape of actin, either because restraining those coordinates traps you in the starting state, or because a MetaD bias can quickly push you into unphysical regions of configuration space. ^40,46^ Other related efforts have investigated the role of flattening on ATP hydrolysis catalyzed by actin, and analogous transitions in the homologous proteins Arp2 and Arp3. ^44,46–50^ None of these previous studies have been able to identified intermediate structures that might occur during flattening.

Here, we report for the first time biased MD simulations that sample reversibly the flat to twisted transition of actin by using our method to produce a position linear discriminant analysis (posLDA) ^32^ coordinate separating the two states. To determine the LDA reaction coordinate, we performed two short MD simulations starting from each of these states and used 10 ns from the twisted and 5ns from the flat state (shorter because it eventually flattens; ^47^ see Sec. A1 for full details). We then performed iterative alignment of all frames in both states (using positions of all 375 C_α_ atoms) to the global mean and covariance as described in. ^32^ LDA on the resulting aligned trajectory yielded a single posLDA coordinate that separates the twisted and flat states. The coefficients for the posLDA coordinate separating the two states is illustrated using a porcupine plot in Fig. 4A. We then performed the OPES variant of WT-MetaD ^51,52^ along this reaction coordinate as described in Sec. A1.

Frame-weighted shapeGMM trained on an OPES MetaD trajectory indicates that five distinct structural states can be occupied during a twisted to flat transition of actin. The trajectory generated contains two full round trip trajectories between flat and twisted states as measured by changes in *ϕ* (Fig. S6), which provides sufficient sampling to investigate the observed conformations and approximate relative free energies. The FES estimated from this approach is shown in Fig. S6. To increase the number of samples available for clustering purposes, we initiated new simulations using a fixed bias taken from the end of the simulation as described in Sec. A1. A cluster scan using these additional frames (see Fig. S7) shows small kinks at *K* = 3 and *K* = 5, and in Fig. 4B,C we show results for *K* = 5 in more detail. Reasonable agreement between the training set and the cross validation set in Fig. S7 demonstrates a lack of overfitting on this data set.

The FES computed from the shapeGMM probability density (*K* = 5) agrees well with the MetaD free energy. Fig. 4B shows the FESs computed from the shapeGMM probability density (in the colormap) and the MetaD (in the contours). The FESs are shown in the space of the *ϕ* and *d* coordinates illustrated in Fig. 4A which have been used to describe the G- to F-actin transition, for better comparison with earlier MD studies. ^40,46^ The MetaD simulation was performed in *ϕ* and the LD coordinate so was reweighted into these coordinates using the same weights fed into shapeGMM. There is impressively good quantitative agreement between the surfaces up to 3 kcal/mol (∼ 5 *k*_B_*T*) considering the very high dimensionality of the GMM. The agreement around the energy minima in this space indicate that the shapeGMM probability density is a good representation of the MetaD simulation results for these regions.

The five states predicted by shapeGMM are in stark contrast to the two that would be predicted just by looking at a 2D free energy projection. Overlain on the FES depicted in Fig. 4B are circles indicating the average *ϕ* and *d* for the structures assigned to each cluster, with the size indicating their relative population. The five state clustering detected two clusters in the flat F-actin like basin (*ϕ* ∼ − 3) and three states in or around the twisted basin (*ϕ* < − 10). The 2D FESs either in *d* and *ϕ* (Fig. 4B) or in the sampled *ϕ* and *LD* (Fig. S6) space have two basins. Clustering in this space would thus likely yield two states. The five-state shapeGMM probability density, however, quantitatively matches the 2D FES thus demonstrating the potential oversimplification achieved in lower dimensional clusterings.

Fig. 4C,D show representative snapshots from the frames assigned to each cluster from the front and above. To give some interpretation to these three different states, we have computed the average root-mean-squared deviation (RMSD) to several published crystal or CryoEM structures of actin alone (twisted), in a filament (flattened), or in complex with an actin binding protein for the C_α_ atoms available in all crystal structures (numbers 7-38, 53-365 out of a total 375). The twisted states (*C* = 0, 1, 4) all had lower RMSD to twisted than flat actin subunits, while the converse is true for the flat states (*C* = 2, 3). State *C* = 4, which is the most twisted, has the lowest RMSD to the starting structure 1NWK (1.67 Å) and ADP-bound actin 1J6Z ^53^ (1.73 Å) than do clusters 0 and 1 (2.59 Å, 2.48 Å). It is expected based on earlier work that our simulations would produce a more flat equilibrium state for ATP-bound actin than what is seen in the crystal structure (which was solved with a non-hydrolyzable ATP analog ^54^). What is interesting is that the clustering algorithm still picks up on this more twisted state as a possible structure, despite the fact that early frames in the trajectory have relatively low weight (since they have little bias applied at that point).

Interestingly, states *C* = 0 and *C* = 1 have equally low RMSD to actin structures in complex with another protein as to the twisted structures considered, for example 2.59 Å and 2.48 Å RMSD to the twisted starting structure 1NWK, but 2.28 Å and 2.09 Å to the structure of actin complexed with the protein profilin (3UB5^55^), which is how a large fraction of actin monomers are found in cells. This suggests that our weighted GMM models may be able to point us towards biologically relevant configurations within a conformational ensemble.

Within the flat states, the most noteworthy difference appears to be in the disordered D-loop (upper right), with cluster 3 having a significantly higher variance than cluster 2. This difference is also evident if we look at the Root-Mean-Squared-Fluctuations of the D-Loop residues shown in Fig. S8. This lower RMSF state (*C* = 2) could correspond to one of the intermediates previously probed through MetaD simulations along a disordered-folded pathway for the D-loop, which were metastable for the ATP-bound actin used in our study, but would be expected to become more stabilized after conversion to ADP. ^56^ Meanwhile, on close inspection (*C* = 3) seems to contain some more disordered structures and some partially folded structures, meaning that the higher variance could be a result of combining two sub-populations into one single state. As it stands, both flattened states have higher RMSF than all twisted states, suggesting a coupling between D-loop structure and twisting that was previously ascribed to nucleotide state (ATP vs ADP), as opposed to the conformational transition which results in ATP hydrolysis, and this would be an interesting question to consider in the future.

## Conclusions

In this work, we present a probabilistic structural clustering protocol that can rigorously account for nonuniform frame weights. This ability allows shapeGMM to be applied, directly, to reweighted or enhanced sampling simulation data to achieve a clustering of in the underlying Hamiltonian of interest. Additionally, we demonstrate that the resulting shapeGMM probability density is a good approximation to the underlying unbiased probability and can thus be used to calculate important Thermodynamic quantities such as relative free energies and configurational entropies. This is a significant advance in our ability to quantify biomolecular ensembles.

By applying our method to actin, we have shown that this approach is capable of picking out physically meaningful structural clusters even for highly complex systems, and illustrates how structural clustering on biased data can provide additional insights that would be difficult to obtain only by looking at the free-energy projected into low dimensional coordinates.

In the future, we envision this approach to be useful in quantifying important biophysical processes such as ligand binding and allosteric regulation.

### A1 Simulation Details

Input files, shapeGMM objects, and analysis codes used to generate all figures are available from a github repository for this article: https://github.com/hocky-research-group/weighted-SGMM-paper

### Beaded Helix

A 12-bead model designed to have two equi-energetic ground states as left- and right-handed helices ^31^ was simulated in LAMMPS. ^57^ 11 harmonic bonds between beads having rest length length 1.0 and spring constant 100 form a polymer backbone. Lennard-Jones (LJ) interactions between every *i, i* + 4 pair of beads withΣ = 1.5 and a cutoff length of 3.0 give rise to the helical shape. The *ϵ* value of this interaction dictates the stability of the helices and was the focus of our reweighting. Simulations were performed with *ϵ* = 6 as the baseline and with *ϵ* = 8 and *ϵ* = 4.5 to assess the accuracy of the reweighting scheme. All non-bonded *i, i* + 2 and farther also have a repulsive WCA interaction with *ϵ* = 3.0 andΣ = 3.0 added to prevent overlap, with the *ϵ* for *i, i* + 2 reduced by 50%. Simulations at temperature 1.0 were performed using ‘fix nvt’ using a simulation timestep of 0.005 and a thermostat timestep of 0.5. A folding/unfolding trajectory of length 50000000 steps was generated and analyzed as above. Here, all parameters are in reduced (LJ) units.

### Alanine dipeptide in vacuum

Alanine dipeptide simulations were performed using GROMACS 2019.6 with PLUMED 2.9.0-dev. GRO-MACS mdp parameter and topology files are obtained from previous PLUMED Tutorials (Belfast-7: Replica Exchange I). AMBER99SB-ILDN force field is used with a time step of 2 fs. NPT ensemble is sampled using velocity rescaling thermostat and Berendsen barostat with a temperature of 300K and pressure 1 bar. For METAD simulations we used PACE=500, SIGMA=0.3Å (for both *ϕ* and *Ψ*) and HEIGHT=1.2 kcal/mol. PLUMED input files are available in our paper’s github repository for complete details.

### Actin monomer

Actin simulations were also performed using GRO-MACS 2019.6 with PLUMED 2.9.0-dev. G-actin with a bound ATP was built and equilibrated at 310 K as described previously. ^47^ The structure of the twisted, ATP-bound actin is derived from the crystal structure with PDB ID 1NWK, ^54^ while that in the flat state is taken from PDB ID 2ZWH, ^58^ with the nucleotide, magnesium ion, and surrounding water replaced with ATP as described previously. MD simulation for ∼ 5ns was performed to relax the starting structure. NPT simulation was performed with 2 fs time step. Parrinello-Rahman barostat is used along with velocity rescaling thermostat with a temperature of 310K and pressure 1 bar. For OPES we used PACE=500, BIASFACTOR=12, BARRIER=15.0 kcal/mol and a multiple time step stride of 2. Two UPPER WALLS were employed ∼ -1^∘^ and 31Å for *ϕ* and *d* respectively. We also used one UPPER WALLS at +40.0 and one LOWER WALLS at -40.0 for the posLDA coordinate. All the walls used were quadratic with a spring constant of KAPPA=500 kcal/mol/nm^2^. PLUMED input files are available in our paper’s github repository.

We performed ∼ 1µs of sampling along this LD coordinate and dihedral angle *ϕ* using the On the Fly Probability Enhanced Sampling variant of Metadynamics (OPES-MetaD). ^51,52^ This method uses a kernel density estimate of the probability distribution over the whole space for biasing rather than building this bias through a sum of Gaussians. The bias at time *t* for CV value ***s***_*i*_ is given by the expression

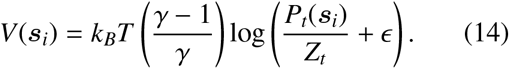

Here, *P*_*t*_(***s***) is the current estimate of the probability distribution, *Z*_*t*_ is a normalization factor. Finally,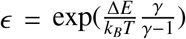 is a regularization constant that ensures the maximum bias that can be applied is Δ*E*. OPES-MetaD data can be reweighted similarly to standard WT-MetaD, using the exponential of the bias (which is similar to rbias for MetaD) or using the estimated free energy of each frame from the final bias. ^51^

We chose the OPES variant of MetaD because (a) literature precedent suggests that it converges more quickly than standard WT-MetaD, and (b) it allows us to set an free-energy cutoff above which bias is not applied (in this case 15 kcal/mol) which limits the amount of unphysical exploration, in a similar manner to Metabasin-MetaD that we previously showed was desirable for this problem. ^46^ Even with this energy cut-off, we needed to include upper and lower walls to prevent over-flattening or over-twisting observed here and in prior attempts by us. ^47^

A cluster scan on our OPES trajectory (Fig. S7) showed a large difference between training and cross-validation curves. Hence we decided to generate to generate additional training frames. We did this by taking the bias accumulated after 900 ns of OPES simulation, and started four 1 ns simulations with random velocities from each of 191 initial configurations from the initial trajectory (separated by 5 ns each), saving every 5 ps; this resulted in ∼ 153k frames available for clustering. The resulting training and cross-validation curves are in much better agreement as discussed in the main text, hence these data were used for clustering and analysis.

## SUPPORTING FIGURES

### S1 Choosing Training Data

When fitting a shapeGMM, we split our data into a training set and a cross validation set. The Gaussian mixture components are fit on the training data and their ability to model the cross validation set is assessed by comparing the log likelihood per frame on both sets. Overfitting will lead to a lower log likelihood on the cross validation set than on the training set. Both training and prediction routines now have built in frame weight arguments.

Training sets were chosen uniformly randomly for the original implementation of shapeGMM. For non-uniform frame weights, however, there are a variety of other methods one could consider to best choose a training set. We assessed a number of these including simple ranking, Poisson sampling, and a Metropolis Monte Carlo method using log frame weights as energies. It was found the the uniform sampling of frame weights worked as well as other methods especially when training sets are sufficiently large.

A uniform sampling of the training set performs at least as well as importance sampling of the training set for the beaded helix example. To assess this we compared shapeGMM objects fit using various training set sampling schemes. These include: a uniform sampling, a Monte Carlo sampling in which frames are replaced based on the Metropolis criteria using frame weights, and a Poisson sampling scheme in which frames are sampled from the frame weight distribution. The Poisson sampling method differs from the other two in that frames are equally weighted in the training set but can appear multiple times depending on their relative weights. The Jensen-Shannon divergence (JSD) between distributions fit using these methods to an *ϵ* = 6 trajectory with *ϵ* = 8 weights and distributions fit to an *ϵ* = 8 simulation directly (the ground-truth; GT) as a function of training set size are depicted in Fig. S1. The JSD between the GT and all fitted distributions is large (∼0.3) for small training set sizes and tends to zero as training sets increase. This indicates that all methods are accurately reproducing the GT distribution for large enough training set. We find that the uniform sampling approach does as well or better than either importance sampling approach for all training set sizes. We note that this result will depend on the specific distribution of weights. We expect this behavior to hold for relatively uniform distributions of weights which occur in reweighting to Hamiltonians that don’t deviate much from the original. It may be important, especially for small training set sizes, in cases in which the Hamiltonians are significantly different to consider choosing training sets using an importance sampling approach. We use a uniform sampling approach for all other applications in this paper.

**Figure S1:**
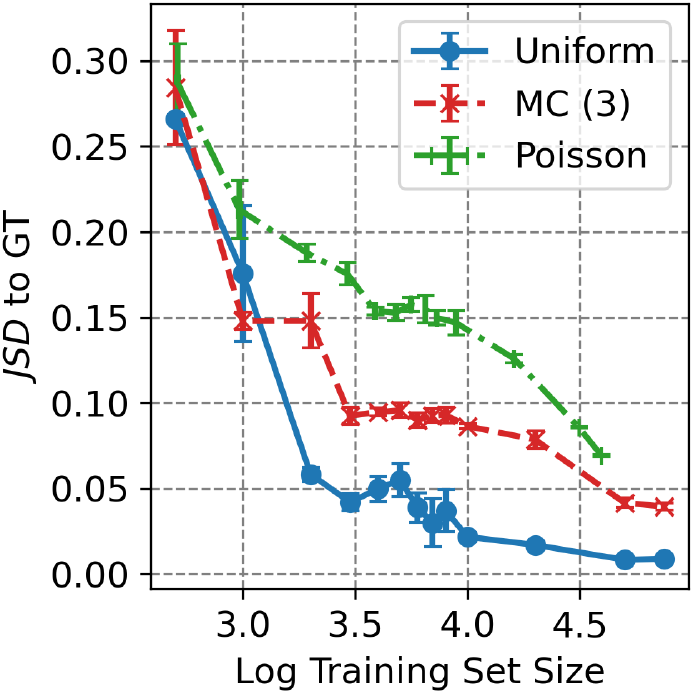
Accuracy of beaded helix reweighted cluster as a function of training set size. The Jensen-Shannon divergence (JSD) between shapeGMM distribution fit using reweighting to *ϵ* = 8 and the ground-truth fit to a simulation run at *ϵ* = 8 as a function of training set size. Three taining set selection schemes are compared: a uniform sampling of frames, a three-step Monte Carlo importance sampling method, and a Poisson sampling method.

### S2 Clustering untempered metadynamics

**Figure S2:**
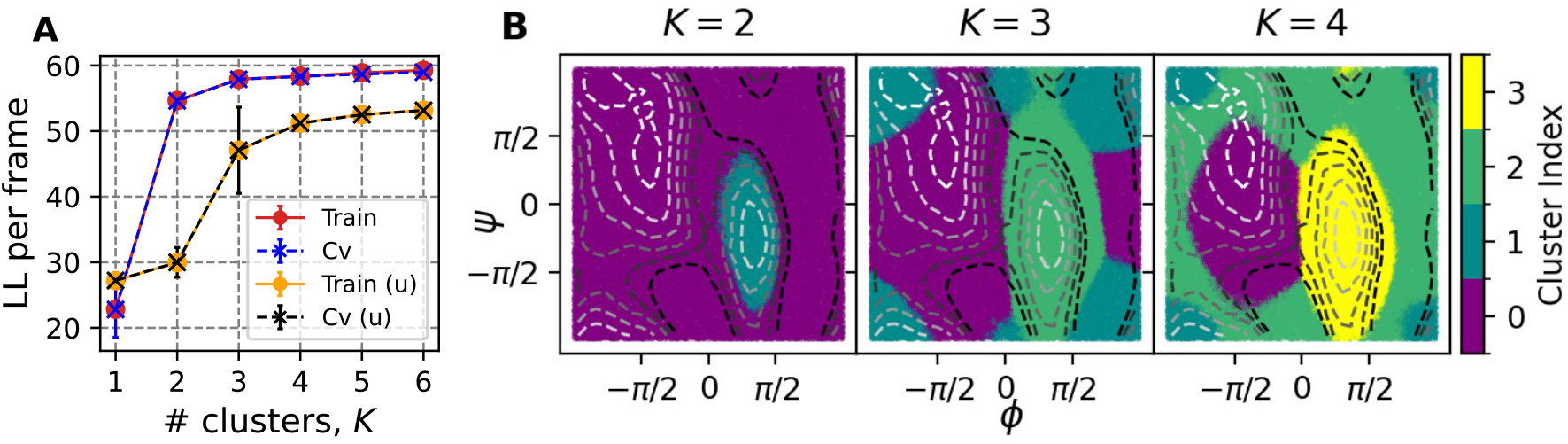
Untemepered MetaD simulation of ADP. (A) Cluster scans obtained with 50K frames, 4 training sets and 10 attempts using rbias frame weights or with uniform weights (labeled ‘u’). Training ln(*L*) curve is substantially higher with rbias weights, and matches CV curve. (B) Clusterings performed for *K* = 2−4 shown by coloring each of 100K sampled points by their cluster assignment. Contour lines indicate the underlying free energy surface as computed from the MetaD simulation via reweighting with rbias frame weights. Contours indicate free energy levels above the minimum from 1 to 11 kcal/mol with a spacing of 2 kcal/mol.

### S3 ADP FES computed by evaluating GMM on WT-MetaD samples

In Fig. S3 we assess an alternative approach to estimate an unbiased FES from a GMM object. In this case, we presume that the WT-MetaD simulation produced physically reasonable configurations spanning the configurational landscape of the molecule of interest. To estimate the FES for ADP, we compute a weighted histogram of (*ϕ* and *Ψ*) where we give as weights the probability of each frame predicted by the GMM, *P*(***x***_*i*_) given by Eq. 2. In practice, *P*(***x***_*i*_) is computed from exponentiating the log-likelihood of frames within the GMM. We normalize the resulting histogram by samples in each bin, which accounts for the fact that frames were not generated uniformly by WT-MetaD, resulting in a new distribution 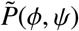. The FES is then computed as 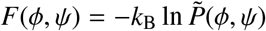

**Figure S3:**
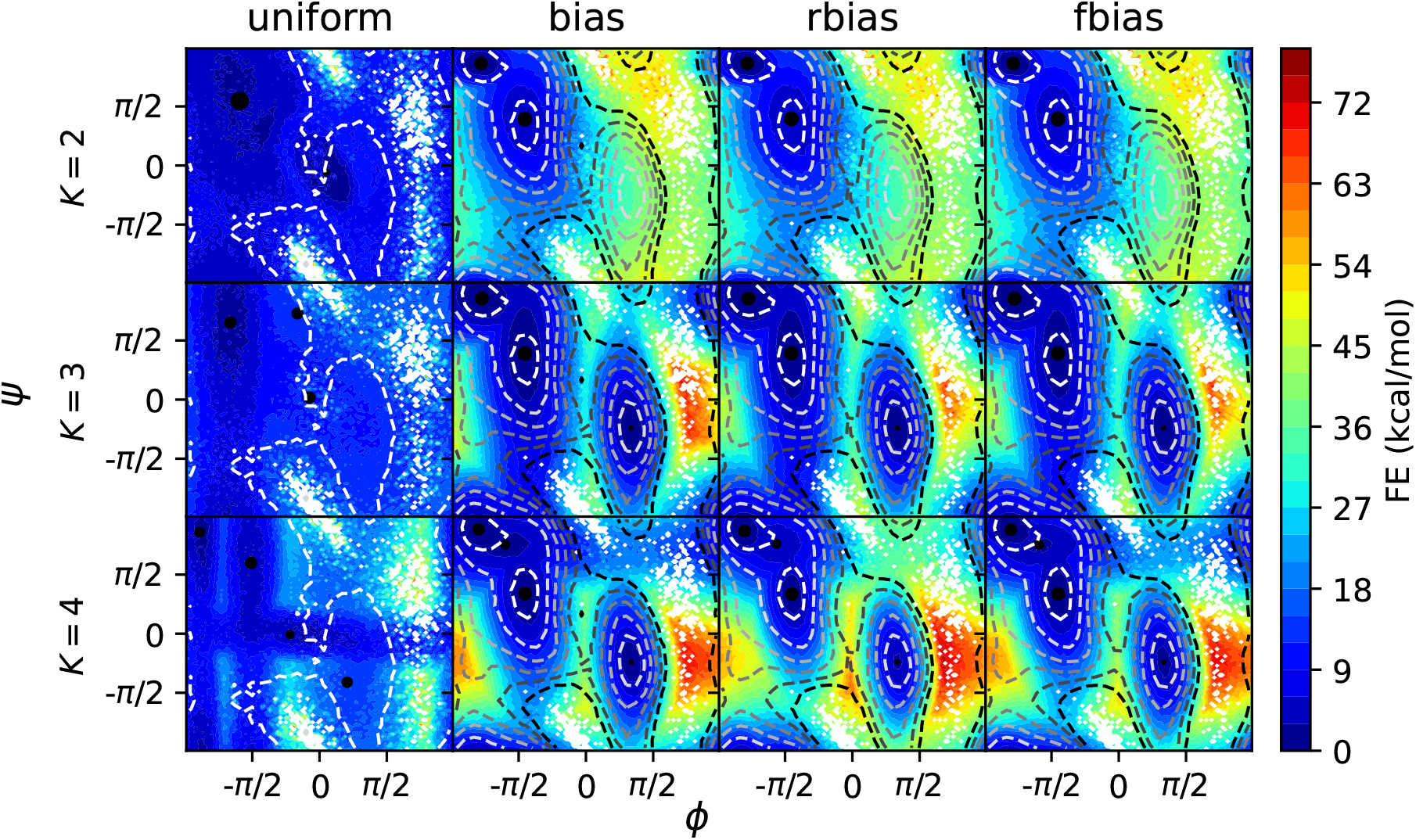
FE profiles obtained from GMM objects trained on BF=10 WT-MetaD data using Monte Carlo procedure. Each column corresponds to a different choice of bias and each row corresponds to a different number of clusters (*K*) used. Black circles placed on the FEs are the centers calculated from the reference structures corresponding to different clusters, with the size indicating their relative population. Contour lines indicate the underlying free energy surface as computed from the WT-MetaD simulation, positioned at 1.0 to 11.0 kcal/mol with a spacing of 2 kcal/mol above the global minimum.

### S4 Error analysis for GMM Free energies

**Figure S4:**
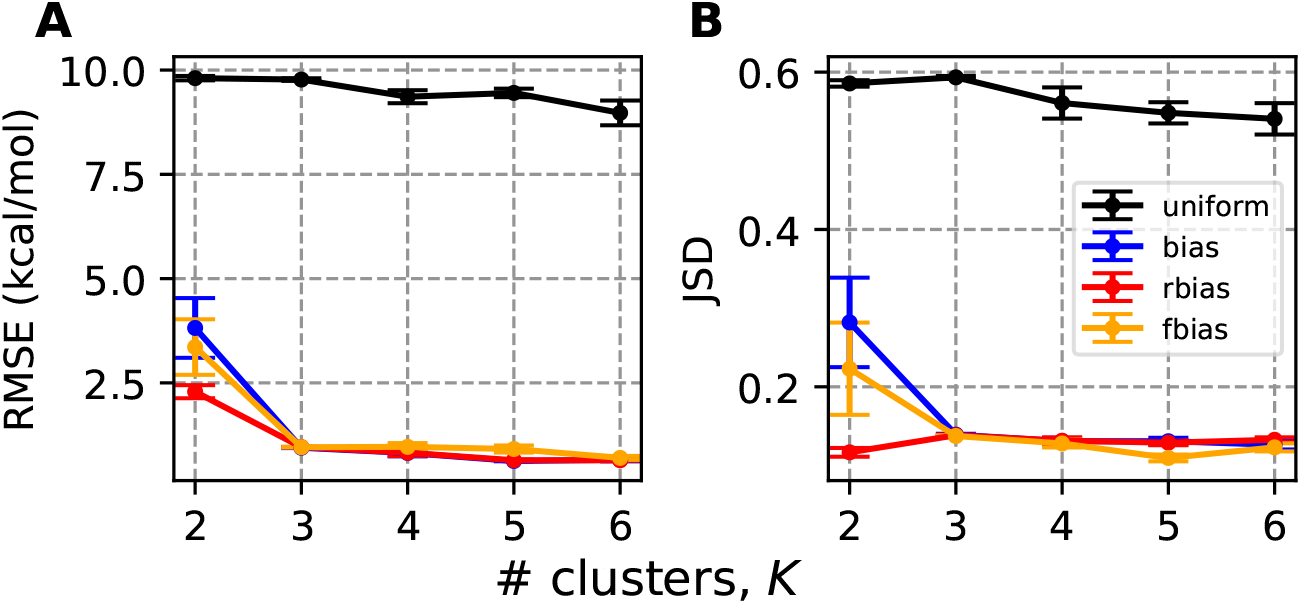
(A) Root mean-squared error for the free energy of ADP GMMs computed for different number of clusters and four different weighting schemes. Error bars are computed from five independent simulations which are fit to separate GMM objects, which are then used to compute free energy surfaces. The reference free energy surface is that computed by summing the Gaussian hills from the WT-MetaD simulation. (B) Same as A, except the Jenson-Shannon distance is computed between the distributions corresponding to *P*(*ϕ, Ψ*) ∝ exp(− *F*(*ϕ, Ψ*)/(*k*_B_*T*)), where *F*(*ϕ, Ψ*) corresponds to either the reference free energy or that computed from the GMM objects.

### S5 FEs from GMM for cluster size 5 and 6

**Figure S5:**
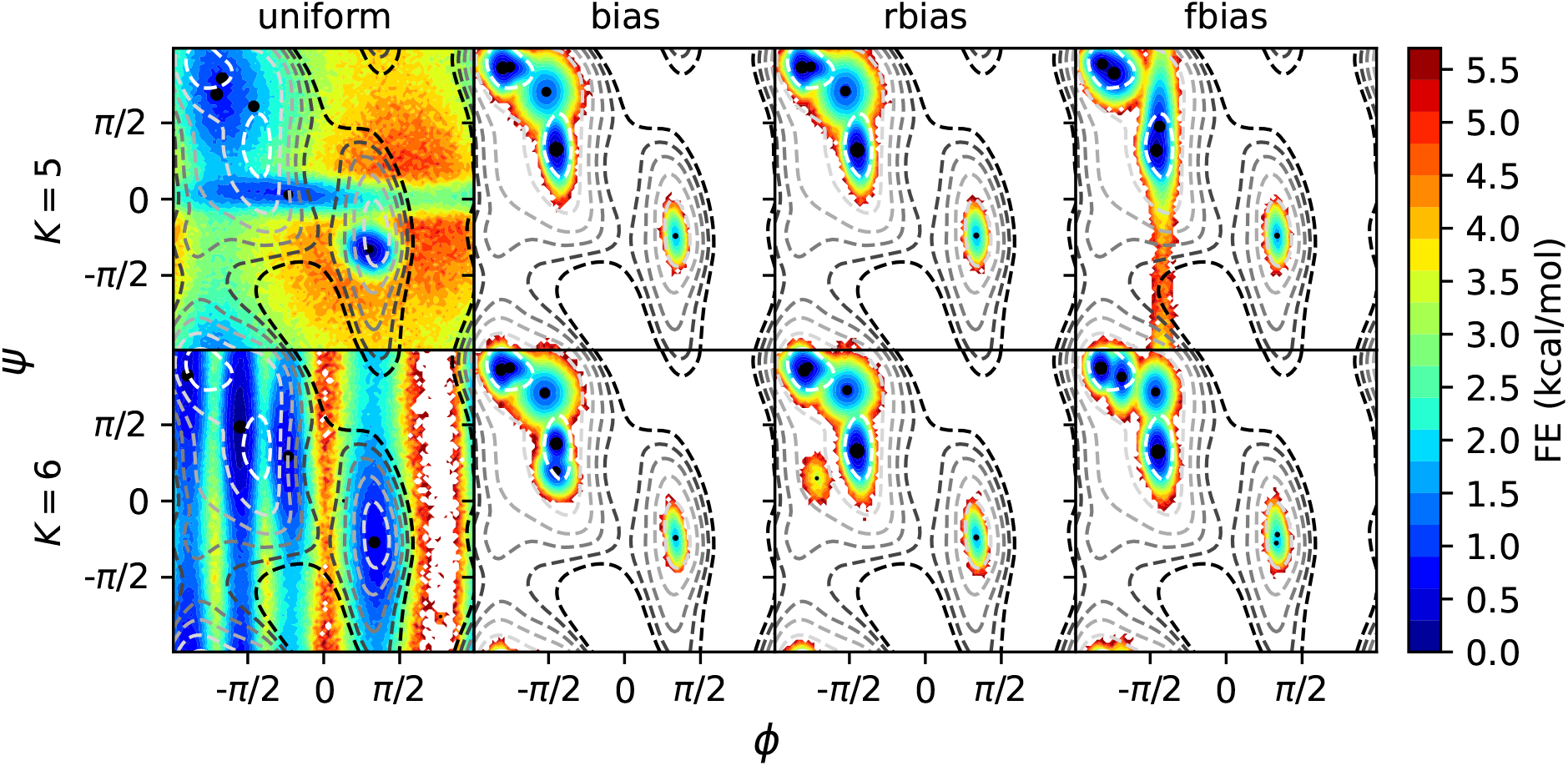
FE profiles obtained from GMM objects trained on BF=10 WT-MetaD data. Each column corresponds to a different choice of bias and each row corresponds to a different number of clusters used. These are computed as unweighted histograms from 1M samples obtained from each GMM object. Black circles placed on the FEs are the centers calculated from the reference structures corresponding to different clusters, with the size indicating their relative population. Contour lines indicate the underlying free energy surface as computed from the WT-MetaD simulation, positioned at 1.0 to 11.0 kcal/mol with a spacing of 2 kcal/mol above the global minimum.

### S6 OPES-MetaD simulation of Actin (∼ 1us)

**Figure S6:**
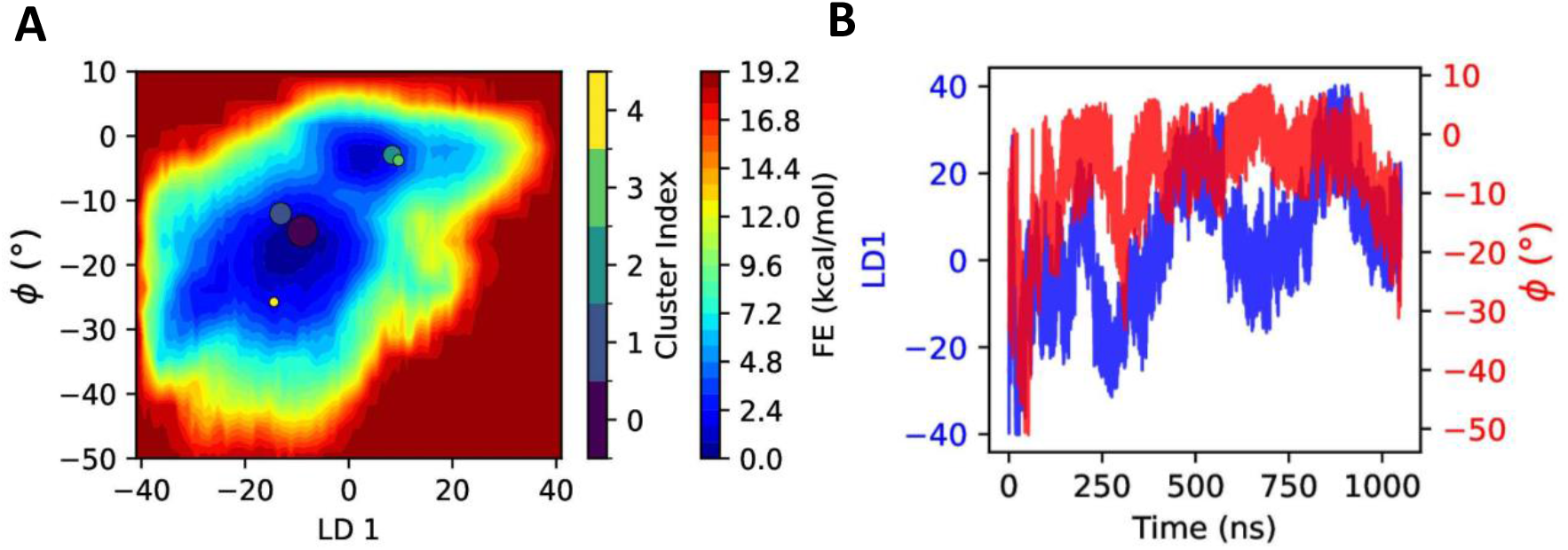
A. The 2D FES obtained from ∼1us OPES-MetaD simulation. Colored circles are the locations for cluster centers weighted according to their relative population. B. Time series of LD1 and Dihedral CVs from the same data.

### S7 Cluster Scans

**Figure S7:**
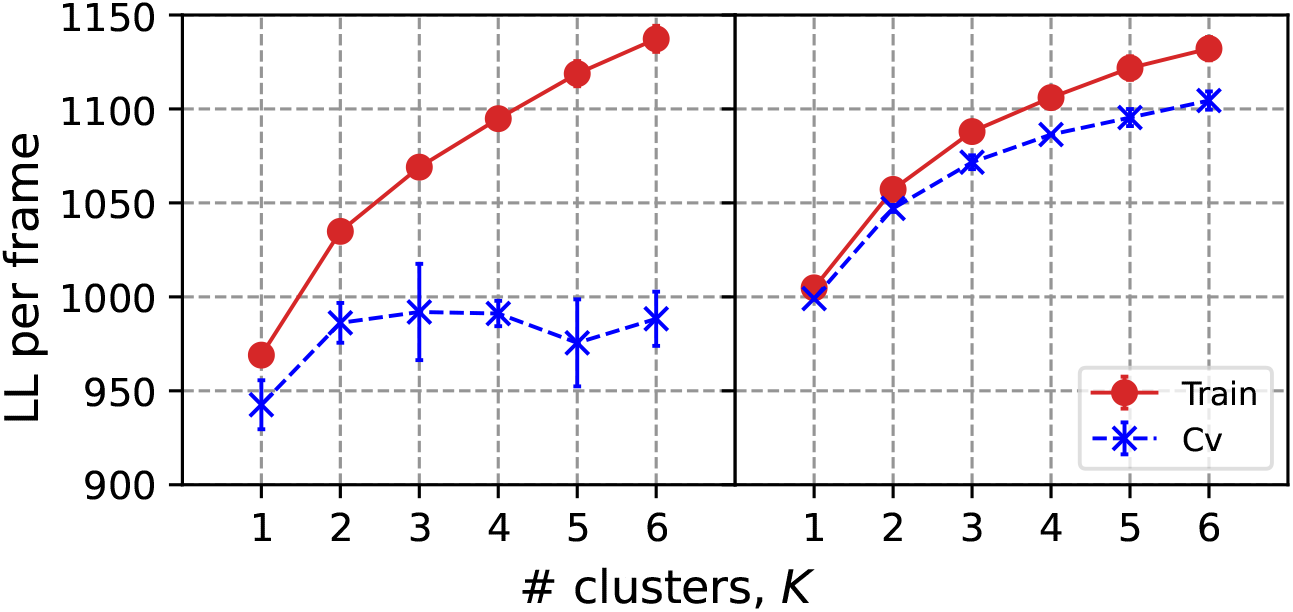
Log likelihood as a function of number of clusters *K* for the original ∼ 1µs OPES-MetaD trajectory (∼21K frames), and using a new set of frames generated by restarting as described in Sec. A1 (∼ 153K frames).

### S8 Variance of D-loop

**Figure S8:**
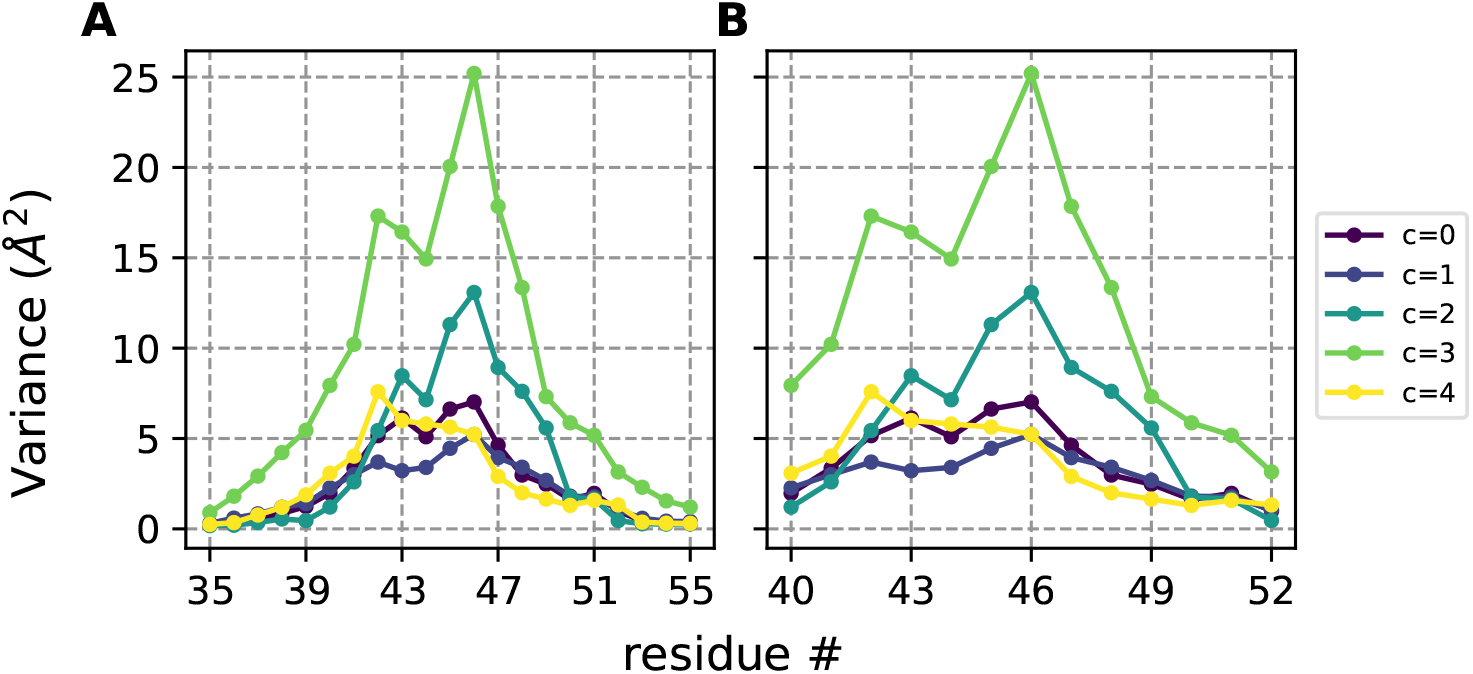
(A) RMSF of Actin’s D-loop for residue 35 to 55 extracted from the diagonal ofΣ_*N*_ for each of five clusters shown in Fig. 4. (B) The same for residues 40 to 52.

